# Crosstalk Between Calcium Dynamics and ROS Levels in U87 Glioblastoma Cells Exposed to Extremely low frequency pulsed electromagnetic fields

**DOI:** 10.64898/2026.04.15.718611

**Authors:** Shahrzad Hadichgeni, Seyed Peyman Shariatpanahi, Bahram Goliaei, Reza Hassan Sajedi, Ali Same-Majandeh, Fahimeh Salehi, Maryam Sadat Nezamtaheri

**Author notes:** Corresponding authors at: P.O. Box 13145-1384, Tehran, Iran. E-mail addresses.

## Abstract

Extremely low-frequency pulsed electromagnetic fields (ELF-PEMFs) have been proposed to modulate intracellular signaling in cancer cells, however, the primary mediators and their temporal sequence remain incompletely understood. In this study, U87 glioblastoma cells were exposed to ELF-PEMF at varying frequencies and amplitudes, and intracellular calcium (Ca^2+^) dynamics, reactive oxygen species (ROS) levels, and mitochondrial membrane potential (ΔΨm) were monitored. Exposure likely led to rapid ROS elevation and a decrease in ΔΨm, indicating early mitochondrial involvement in ELF-PEMF responses. Fast Fourier transform (FFT) analysis of Ca^2+^ oscillations suggested that low-frequency exposures produced higher spectral power and amplitude compared with controls, consistent with enhanced Ca^2+^ signaling activity. Parallel pharmacological experiments demonstrated that ROS elevation may occur independently of IPL-dependent endoplasmic reticulum (ER) Ca^2+^ release, as 2-Aminoethoxydiphenyl borate (2-APB) inhibition did not prevent ROS increase. In contrast, treatment with the antioxidant N-acetylcysteine (NAC) effectively suppressed ROS without significantly altering basal cytosolic Ca^2+^ levels. These observations indicate that ROS likely acts as an early mediator of cellular responses to ELF-PEMF exposure, with downstream modulation of calcium signaling pathways. The magnitude of ROS elevation and Ca^2+^ modulation was strongly dependent on field frequency and amplitude, consistent with a frequency-dependent biological window. Overall, ROS likely acts as a primary mediator of ELF-PEMF bioeffects, highlighting its potential relevance for glioblastoma therapy, and future studies are warranted to assess other glioma lines to confirm generalizability.

## 1. In**troduction**

Extremely low-frequency pulsed electromagnetic fields (ELF PEMFs) are pervasive in modern life, yet their biological effects remain incompletely understood. Traditional studies have primarily focused on calcium signaling and mitochondrial membrane potential (MMP) as key targets. Emerging evidence suggests that reactive oxygen species (ROS) may act as central upstream mediators, orchestrating downstream changes in Ca²L dynamics and mitochondrial function, and can rapidly elevate intracellular ROS via cryptochrome and mitochondrial activation, even before significant alterations in Ca²L influx or MMP occur (1, 2). Exposure to ELF-PEMFs has been shown to increase ROS levels, leading to oxidative stress. This process is accompanied by alterations in antioxidant enzyme activity and elevated levels of oxidative damage markers, including malondialdehyde (MDA) and protein carbonyls (3). In muscle cell models (e.g., C2C12 myoblasts), ELF PEMF induces ROS accumulation, ΔΨm reduction, and changes in intracellular Ca²L handling (4). Mitochondria, as the main ROS source, regulate both Ca²L homeostasis and ΔΨm, forming a bidirectional feedback circuit between redox status and calcium signaling (5–7). Local Ca²L transfer to mitochondria occurs via IP3 receptors (IP3R), stimulating mitochondrial oxidative metabolism. This process is independent of outer mitochondrial membrane channels and allows dynamic Ca²L shuttling between ER–mitochondria contact sites (ERMCs), supporting local communication(8). Mitochondrial ROS can modulate IP3R activity and shape Ca²L oscillations, while Ca²L fluctuations reciprocally influence ROS production, forming a tightly coupled regulatory network ((9)). Calcium acts as a versatile intracellular signal across temporal and spatial scales. Rapid, localized transients (calcium flashes) elicit immediate responses such as exocytosis, mitochondrial redox regulation, and gene transcription, interpreted through frequency encoding. Global or repetitive calcium waves coordinate longer-term responses, whereas sustained high intracellular Ca²L ([Ca²L] i) is generally associated with pathological outcomes, including cell death, mitochondrial overload, and ΔΨm collapse(10–12). Spectral analyses, including FFT, can be used to convert calcium signals from the time domain to the frequency domain and quantify the oscillatory components of Ca²L(13). Despite these insights, the causal hierarchy between ROS, Ca²L, and ΔΨm under ELF PEMF exposure remains unclear (14). It is unknown whether ROS acts upstream via IP3R to drive calcium oscillations, or if calcium dynamics precede ROS production. This is particularly relevant for glioblastoma, where IP3R3-mediated calcium signaling promotes proliferation, migration, and invasiveness. Elevated mitochondrial Ca²L and ROS may trigger mitochondrial permeability transition pore (mPTP) opening, leading to apoptosis; yet, cancer cells suppress cell death pathways through regulation of calcium channel assembly and ROS scavenging (15–17). In this study, we aim to clarify the hierarchical interaction between ROS and Ca²L under weak ELF PEMF exposure in glioblastoma cells. By simultaneously monitoring ROS, intracellular calcium ([Ca²L]i), and ΔΨm, and employing pharmacological interventions such as 2-APB (IP3R inhibitor) and N-acetyl-L-cysteine (ROS scavenger), we investigate whether ROS acts as the upstream driver of calcium dynamics or if calcium changes precede ROS production. Understanding this interplay will provide both biophysical insights into ELF PEMF bioeffects and mechanistic insights into the intracellular signaling pathways, and may inform potential therapeutic strategies for diseases in which oxidative stress and calcium signaling are pathophysiologically significant.

## 2 Materials and Methods

### 2.1. Cell Culture

Human U87 glioblastoma cells (ATCC, USA) were maintained in Dulbecco’s Modified Eagle Medium/Nutrient Mixture F-12 (DMEM/F-12; Gibco, UK) supplemented with 10% fetal bovine serum (Gibco), 100 IU mlL¹ penicillin, and 0.1 mg mlL¹ streptomycin (Sigma-Aldrich). Cells were acquired from the Iranian Biological Resource Center (IBRC, Iran). Cultures were incubated at 37 °C in a humidified atmosphere containing 5% COL (Heraeus, D6450 Hanau), and all experiments were performed using cells in logarithmic growth phase. Cell viability was quantified using the Trypan Blue exclusion assay and counts were obtained using a hemocytometer.

### 2.2. ELF-PEMF Exposure

Cells were exposed to ELF-PEMF using a custom-designed exposure system as described in previous studies(2, 18). The system consists of an excitation coil wound around a magnetic core, creating a uniform magnetic field within an air gap where culture dishes are placed. This design ensures homogeneous field distribution, minimizes thermal artifacts, and allows precise control of field intensity and pulse timing. In this study, cells were exposed to a train pulsed magnetic field with a magnetic flux density of 100 mT and periods of 7, 20, 70, and 200 seconds for a total duration of 45 minutes. The pulse width was set to one-fifth of the total period (0.2 duty cycle). For example, a pulse train with a 20-s period had a pulse width of 4 s. Under all experimental conditions, these values refer to the pulse period, while the pulse width corresponded to either one-quarter or one-fifth of each period, depending on the specific condition, with the magnetic field remaining off for the remainder of the cycle. Sham controls were maintained under identical incubator conditions without exposure to the magnetic field. Field homogeneity was verified, and the system was designed to prevent heating or mechanical disturbances, ensuring that any observed biological effects were primarily attributable to exposure to the ELF-PEMF.

### 2.3. ROS Measurement

Intracellular ROS production was quantified using 2′,7′-dichlorofluorescein diacetate (DCFH-DA; 10 µM; Sigma-Aldrich). Following ELF-PEMF exposure, cells were incubated with DCFH-DA for 30 min at 37 °C in phenol-red-free, serum-free DMEM. Because oxidized DCF rapidly redistributes into the cytosol, imaging was performed immediately after dye loading. Fluorescence was measured using Zeiss and Nexcope NIB620FL microscopes (FL20 LED module) and quantified using a Synergy H4 plate reader (Ex/Em: 485/535 nm).

### 2.4. Measurement of Intracellular Calcium Dynamics

U87 glioblastoma cells (1 × 10L per well) were seeded into 96-well plates and allowed to adhere overnight. Intracellular Ca²L levels were measured using Fura-2 AM (2.5 µM; Sigma-Aldrich). Cells were loaded with the dye for 45 min at 37 °C in the dark in HBSS buffer (137 mM NaCl, 5.4 mM KCl, 0.34 mM NaLHPOL, 1 mM MgCl, 10 mM glucose, 0.4 mM MgSO_4_-7H_2_O, 0.44 mM KH_2_PO_4_, 5 mM CaClL, ≈12 mM HEPES, pH 7.4) supplemented with 1 mg mlL¹ BSA and 0.01% Pluronic F-127(19, 20). After loading, cells were washed and incubated in dye-free HBSS for an additional 45 min to allow complete intracellular de-esterification. During the second 45-minute interval, cells were exposed to ELF PEMF (100 mT) using a custom-designed exposure system under controlled temperature and COL conditions and a segmented temporal stimulation pattern consisting of four discrete pulsed periods of 7, 20, 70, and 200 s, corresponding to fundamental frequencies of 0.142, 0.05, 0.0142, and 0.005 Hz. Live-cell calcium imaging was performed immediately after exposure and also one-hour post-exposure to evaluate delayed effects and assess changes in intracellular Ca²L dynamics over time following ELF-PEMF stimulation. Chemical depolarization was induced using standard KCl solutions according to Abcam protocols, with a 50 mM KCl pulse serving as a positive control. Intracellular Ca²L dynamics were measured using time-lapse imaging with Fura-2 on an inverted phase-contrast microscope (Zeiss M405, Germany) equipped with a dual-excitation LED system (F340/F380 nm) and a high-speed scientific camera, with fluorescence emission collected at 510 nm. Images were acquired with a fixed exposure of 10 ms and recorded at variable frame intervals ranging from 0.14 to 0.6 s for up to 2 min. Parallel fluorescence measurements were obtained using a Synergy H4 multimode plate reader (BioTek) to monitor the F340/F380 ratio.

#### 2.4.1. Image Processing

Reactive cells exhibiting dynamic Ca²L responses were identified by manually delineating regions of interest (ROIs) encompassing the entire cell body in Fiji (ImageJ). Prior to quantification, excitation and emission channels were separated, and fluorescence signals were extracted using appropriate Fiji plugins, including the Time Series and Calcium Imaging tools. All image stacks underwent constant background subtraction, and when necessary, a median filter was applied to reduce high-frequency noise prior to ROI-based fluorescence analysis. Exposure settings, thresholding parameters, and all user-defined preprocessing steps were kept identical across experimental groups to ensure comparability and minimize analytical bias. Following background correction, fluorescence intensity was extracted for each cell across the entire imaging sequence, capturing cell-wide changes in signal intensity over time. Baseline fluorescence (FL) was determined from the initial pre-stimulation frames, and all traces were normalized to ΔF/FL, where ΔF/FL = (F − FL)/FL. To quantitatively assess the temporal dynamics of intracellular Ca²L signaling, the variance of the fluorescence intensity over time was calculated for each individual cell. Variance was selected as an unbiased metric to capture the magnitude and temporal variability of Ca²L-dependent fluorescence changes, independent of absolute signal amplitude. For each experimental condition, the mean variance across all analyzed cells (n ≥ 60) was reported as a single quantitative measure. ROI selection criteria, normalization procedures, and analysis parameters were applied consistently across all datasets to ensure methodological reproducibility.

#### 2.4.2. FFT Analysis of Ca²D Oscillations

To resolve the frequency and amplitude components of intracellular Ca²L signals, fluorescence time-series data were normalized prior to fast Fourier transform (FFT) analysis. FFT was computed using custom-written Python scripts. Each of the four datasets was acquired under stimulation with distinct oscillation periods (different frequencies). Therefore, results were plotted separately in four individual graphs rather than combined. The dataset was analyzed in comparison with negative and positive controls.

### 2.5. Dynamic Inhibitor Analysis of Ca²D and ROS Up/Downstream

2-Aminoethoxydiphenyl borate (2-APB; EMD Millipore, USA) was used as a pharmacological suppressor of calcium signaling, as it inhibits store-operated calcium entry (SOCE) and IPL-mediated Ca²L release from endoplasmic reticulum stores (21, 22)(23, 24). A 2 mM stock solution was prepared in DMSO. It was diluted to a final working concentration of 50 µM in experimental buffer, and cells incubated for 1 h prior to Ca²L imaging. N-acetylcysteine (NAC; Sigma-Aldrich), which increases intracellular glutathione availability and effectively scavenges ROS to reduce oxidative stress (23), was used at an optimal concentration of 2.5 mM for 1 h prior to ROS analysis based on preliminary tests assessing ROS reduction and cell viability. Considering that IPL receptors (IPLRs) are functionally coupled to mitochondria, the primary source of intracellular ROS, these two inhibitors (2-APB and NAC) were used to evaluate the relative contributions of cytosolic Ca²L dynamics and ROS in U87 cell responses to ELF-PEMF. Among four predefined stimulation periods (described in Section 2.2), the period exhibiting the most prominent modulation of cytosolic Ca²L dynamics relative to the positive control (50 mM KCl) was selected for further analysis. Experiments were designed to measure cytosolic Ca²L dynamics and ROS levels in parallel for each inhibitor. This approach allowed us to determine which signal (Ca²L or ROS) acts upstream or downstream when U87 cells are exposed to ELF-PEMF at the specified period.

### 2.6. Mitochondrial **ΔΨ**m Evaluation

Mitochondrial membrane potential (ΔΨm) was assessed using tetramethyl rhodamine ethyl ester (TMRE; 100 nM). Cells were incubated with TMRE for 45 min, washed, and imaged in phenol-red-free DMEM. TMRE fluorescence was excited at 549 nm, and emission was collected in the 575–610 nm range using the same Zeiss inverted microscope described above.

### 2.7. Apoptosis Assay

Apoptotic cell death was evaluated using a dual approach combining acridine orange (AO)/ethidium bromide (EB) staining and flow cytometry. For morphological assessment, cells were incubated with AO at 1 µg/mL and EB at 0.5 µg/mL (Sigma-Aldrich), and fluorescence images were acquired to distinguish viable, early apoptotic, and late apoptotic/necrotic cells based on differential staining patterns. For quantitative analysis, flow cytometry was performed using Annexin V-FITC (Catalog No. K101-100; BioVision, USA) in combination with propidium iodide (PI) at a final concentration of 5 µg/mL. This strategy allowed simultaneous evaluation of apoptotic populations both morphologically and quantitatively, ensuring robust and reproducible measurements.

### 2.8. Statistical Analysis

All experiments were independently repeated at least three times. Data are presented as mean ± SEM. For comparisons involving multiple groups, one-way ANOVA followed by Tukey’s post-hoc test was used. A p-value < 0.05 was considered statistically significant. Fluorescence-based measurements, including normalized Ca²L signal intensities, were analyzed using standard high-impact imaging procedures. Signal intensities were normalized to baseline values (F/FL or ΔF/FL) to allow cross-group comparisons. For specific analyses, including ROS quantification and membrane-potential measurements, raw fluorescence values were reported to preserve the original signal distribution. Statistical analyses and graphing were performed using GraphPad Prism 9 (GraphPad Software, USA) and Microsoft Excel, the flow cytometry data were analyzed using FlowJo v11 (BD Biosciences, USA).

### 2.8. Supplementary Movies

Representative time-lapse recordings illustrating key dynamic responses, including Ca²L flashes, ROS fluctuations, and mitochondrial membrane-potential changes (TMRE, ΔΨm), are provided as Supplementary Movies 1–11. These videos correspond to the imaging datasets used for quantitative analyses and are included to allow visualization of the real-time cellular events underlying the reported measurements.

## 3 Results

### 3.1. ELF-PEMF exposure analysis and optimal frequency selection 3-1-1. Cell Intracellular calcium-intensity dynamics

Quantitative analysis of the calcium imaging data revealed that all four groups exposed to ELF-PEMF at 100 mT with periods of 7, 20, 70, and 200 seconds exhibited a sustained increase in the mean variance of intracellular relative calcium fluorescence compared to the negative control. Comparative analysis indicated that stimulation at 20 and 7-second period produced the largest increases in the variance of relative calcium fluorescence, with statistically significant enhancements (P < 0.001) relative to the negative control. The 20-second period (0.05Hz) exceeded the depolarization threshold induced by 50 mM KCl (positive control), showing a greater increase compared to both KCl-induced signals and the negative control. The other two periods (70 and 200 seconds) induced smaller increases in calcium fluorescence variance and did not produce statistically significant changes relative to the negative control. One hour after field exposure, the variance of relative calcium fluorescence in all groups decreased compared to initial recordings; however, none of these changes reached statistical significance relative to baseline. Cellular responses of U87 glioblastoma cells to four pulsed ELF-PEMF periods at 100 mT are summarized in Figure 2A, with significant and non-significant changes indicated relative to the negative control. Figure 2B shows a short non-linear line plot connecting the four stimulation periods, with the highest response observed at the 20-second period. Figure 2C shows the time-series calcium fluorescence signal at the 20-second period, illustrating the temporal up-and-down fluctuations of calcium flash intensities relative to the negative control.

**Fig. 1.**
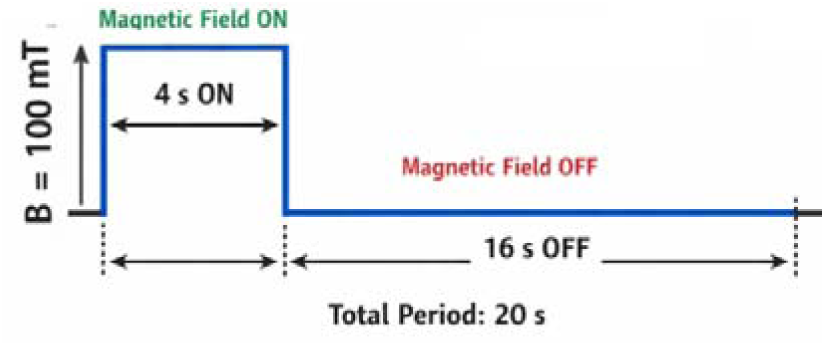
Example of a pulsed magnetic field waveform with an intensity of 100 mT, a 20-s period, and a pulse width of 4 s.

**Fig. 2.**
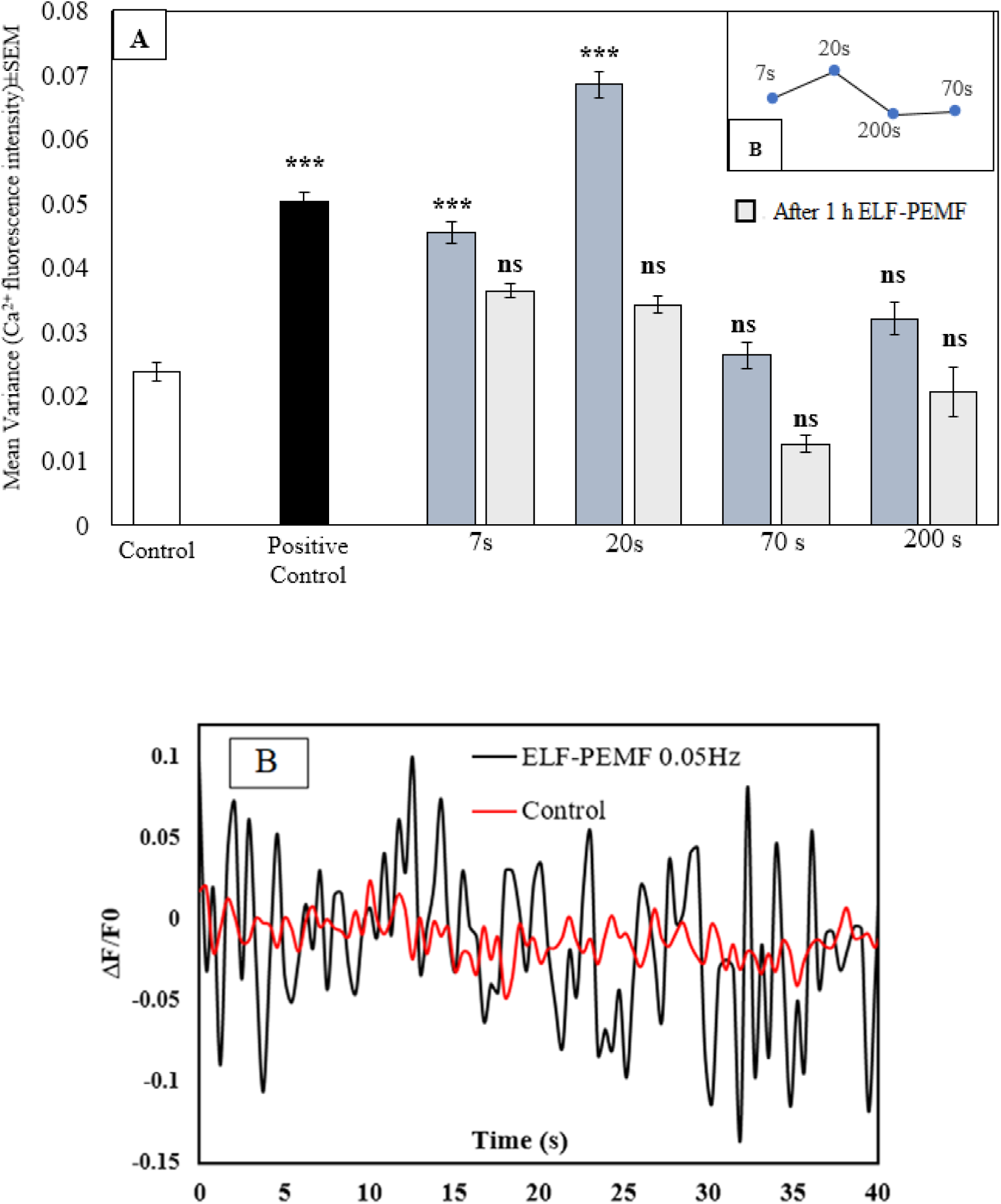
(A) Variance of relative Ca²ϑ fluorescence in U87 cells during 45-min exposure to pulsed ELF-PEMF at four periods (T = 7, 20, 70, and 200 s) at 100 mT. (B) Short non-linear line plot of the four periods, with the highest response at 20 s. (C) Time-series of calcium flashes at the 20-second period, showing up-and-down fluctuations. Data are mean ± SEM. “ns” indicates not significant; *** indicates statistically significant relative to the negative control

#### 3.1.2. FFT-based analysis of intracellular Ca²D oscillation

FFT analysis of intracellular calcium time-series data from cells exposed to ELF-PEMF with four different stimulation cycle durations revealed that the highest oscillation amplitude and spectral power, relative to the negative control, corresponded to stimulation periods of T = 20 (0.05Hz), 7 (0.142 Hz), 200 (0.005 Hz), and 70 (0.0142 Hz) seconds, respectively. This increase in spectral power was predominantly observed at low frequencies below 0.4 Hz, compared with the positive control. Stimulation cycle durations of 20 and 7 seconds also exhibited increased oscillation amplitude and spectral energy, with the difference being most pronounced at the 20-second cycle. In contrast, the negative control (unstimulated cell) displayed lower spectral power amplitudes in the low-frequency range (<0.1 Hz), characterized by a gradual decay consistent with the typical biological 1/f pattern, relative to the ELF-PEMF–exposed groups. Conversely, cells exposed to ELF-PEMF showed elevated spectral power and larger oscillation amplitudes at low frequencies and maintained higher power levels even with increasing frequency compared to controls. Notably, the 20-second stimulation cycle (corresponding to a stimulation frequency of 0.05 Hz) induced the largest oscillation amplitude, the strongest dynamic calcium response, and the highest variance in relative calcium intensity in U87 glioblastoma cells, surpassing both negative and positive controls as well as other stimulation cycle durations and frequencies. Based on these observations, all subsequent experimental assessments in this study were conducted using the 0.05 Hz. Figure 3 shows the FFT spectra of Ca²L fluorescence in U87 cells, with comparisons to both negative and positive controls.

**Fig. 3.**
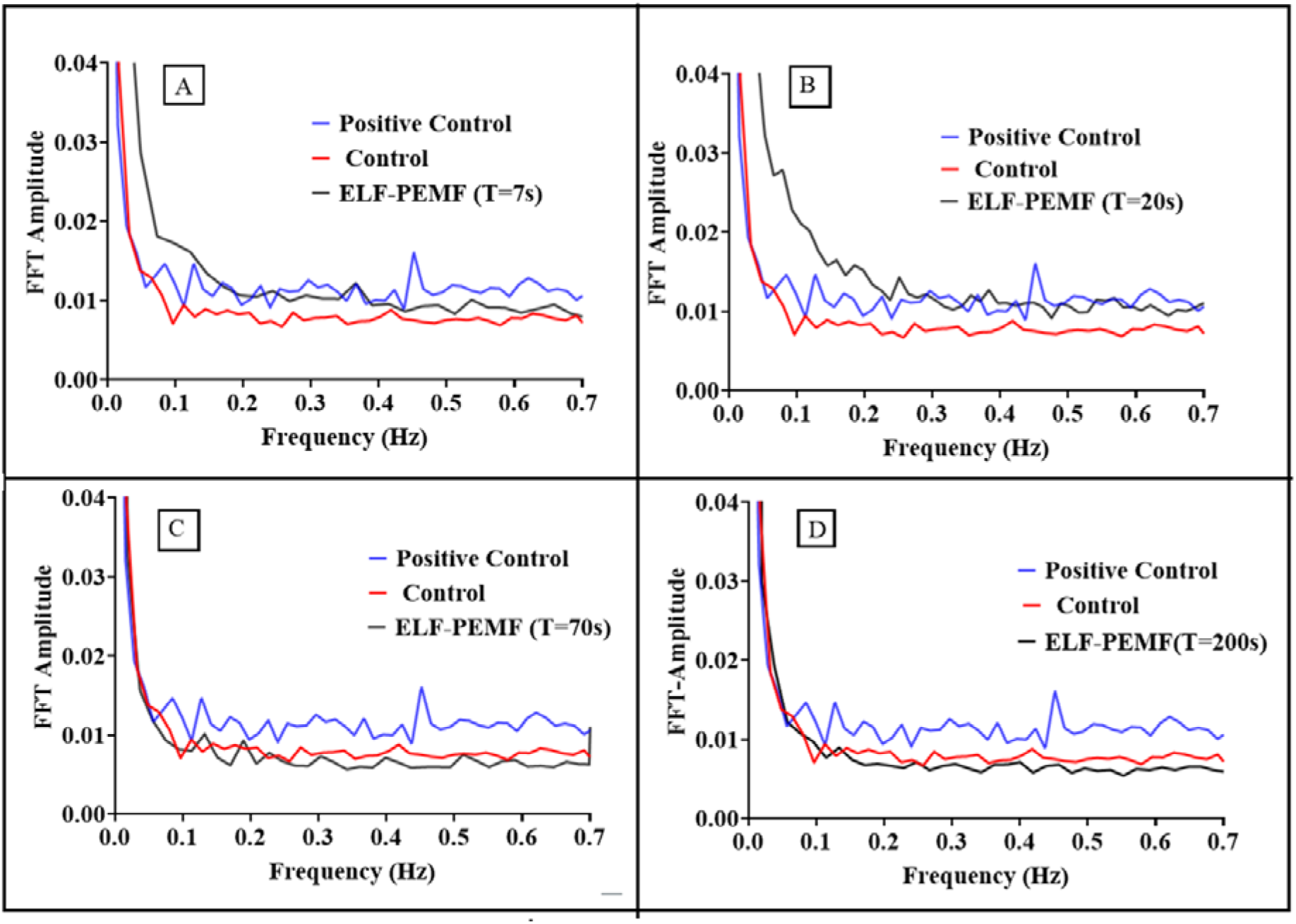
FFT spectra of Ca² fluorescence fluctuations in U87 cells during 45-min exposed to ELF-PEMF stimulation with periods of T= 7 s (A), 20 s (B), 70 s (C), and 200 s (D), at 100 mT, alongside the negative control(unstimulated) and the positive control (50 mM KCl).

### 3.2. Modulation of ΔΨm by ELF-PEMF stimulation at 0.05 Hz

The mitochondrial membrane potential (ΔΨm) was quantified using the fluorescent probe TMRM following exposure of U87 cells to ELF-PEMF for 45 minutes. As shown in Figure 4A, data are shown for comparative analysis of field-exposed versus control cells, the fluorescence intensity of cells under the magnetic field exhibited a gradual reduction, deviating from the linear fluorescence trend observed in negative control cells. Interestingly, the decline followed an approximately stepwise and oscillatory pattern, with apparent steps occurring roughly every 20 seconds, corresponding to the pulse period of the magnetic field (0.05 Hz), as indicated by red arrows in Figure 4B. This rhythmic, step-like depolarization persisted throughout the imaging session. In negative control U87 cells, fluorescence intensity remained largely stable, without the distinct periodicity or amplitude observed in cells exposed to pulsed magnetic fields.

**Fig. 4.**
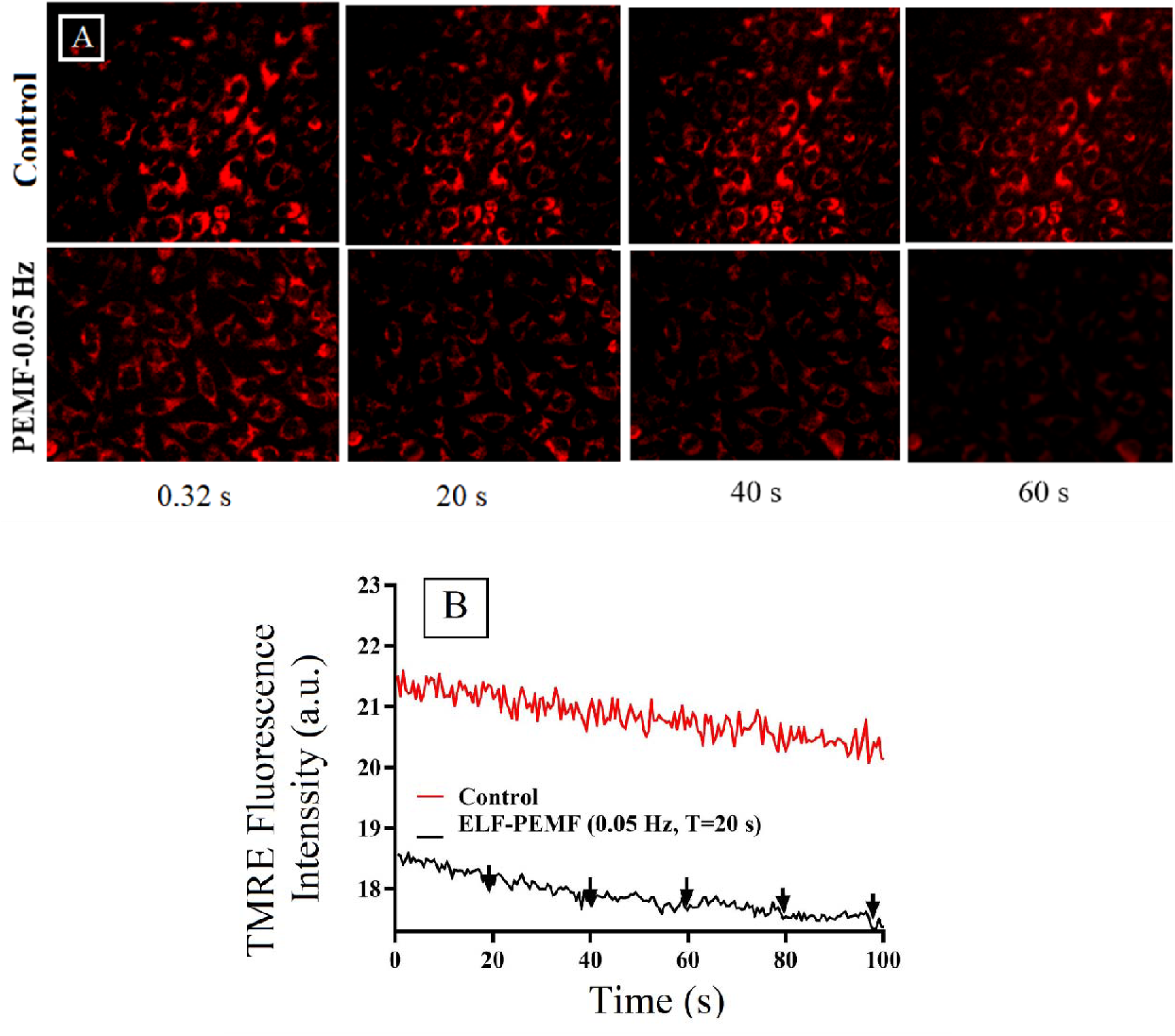
(A) TMRM fluorescence intensity in U87 cells at four time points (0.32, 20, 40, and 60 s) under ELF-PEMF exposure and in the negative control. (B) Time course of TMRM fluorescence under ELF-PEMF exposure. Black arrows indicate stepwise depolarization events corresponding to the magnetic pulses. Images were acquired using an inverted Zeiss fluorescence microscope at 32× magnification.

### 3.3. ROS dynamics

Intracellular ROS dynamics were quantified in U87 glioblastoma cells using a fluorescence-based assay under control conditions and following exposure to an ELF-PEMF (0.05 Hz). ELF-PEMF stimulation induced a marked increase in ROS production compared with controls (Figure 5A). The fluorescence trajectory exhibited a rapid initial rise, reaching a maximum rate of increase of approximately 0.127 a.u.·sL¹ around 15 s, followed by a clear plateau phase and a subsequent gradual decline. In contrast, control cells showed only a modest and transient increase with a slower rate of change (≈0.077 a.u.·sL¹) and no evident plateau. ELF-PEMF–treated cells reached half-maximal ROS levels earlier than controls (tL/L ≈ 24 s vs. ≈ 45 s, respectively (Figure 5B).

**Fig. 5.**
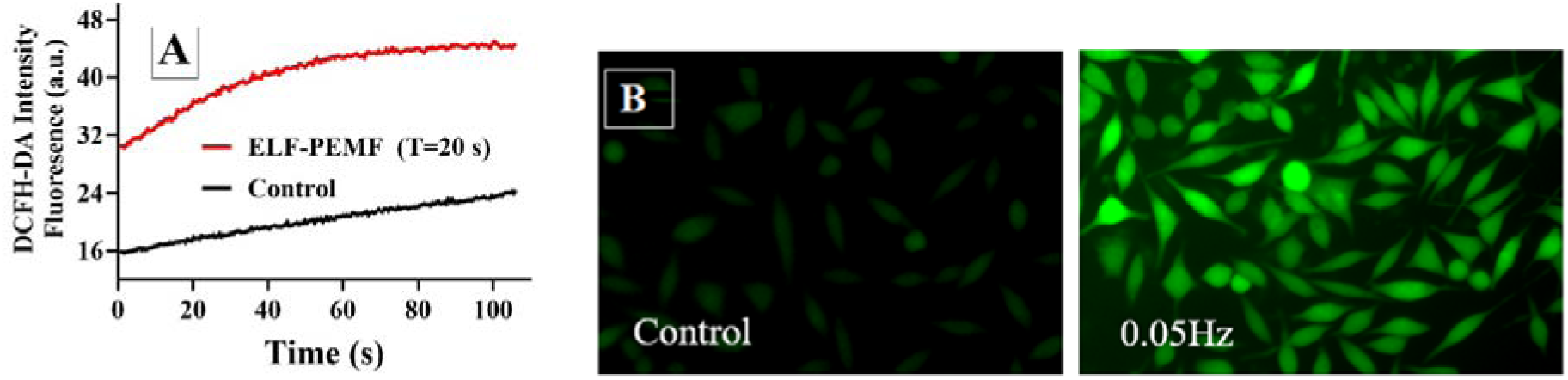
Representative ROS fluorescence images measured with DCFH-DA under 0.05-Hz ELF-PEMF exposure at 100 mT for 45 min, alongside the negative control (A), with quantification shown in (B). Images were acquired using a fluorescence microscope at 40× magnification.

### 3.4. ELF-PEMF Cellular Responses: Upstream of ROS or Ca2+ 3-4-1. NAC and 2-APB Effects on ROS

Treatment of U87 cells with 2.5 mM NAC (a pharmacological ROS inhibitor) under basal conditions resulted in a noticeable reduction in ROS-associated fluorescence intensity. However, when the cells were exposed to a 0.05-Hz ELF-PEMF field and simultaneously treated with NAC, ROS levels showed a slight increase that was not statistically significant compared with the control. Although previous experiments demonstrated that the field alone at this frequency range induces a significant increase in ROS levels, under combined ELF-PEMF and NAC treatment, no significant difference from the control values was observed. While examining ROS levels in U87 cells treated with 50 μM 2-APB, only a slight, non-significant increase was observed compared to the control, indicating that inhibition of Ca²L signaling alone at this concentration does not appreciably affect basal ROS production. However, when cells were simultaneously exposed to the 0.02-Hz ELF-PEMF field and 50 μM 2-APB, ROS levels were significantly higher than in cells exposed to the field alone. These results suggest a potential synergistic interaction between ELF-PEMF stimulation and Ca²L signaling inhibition at this concentration, whereby the combined treatment amplifies ROS accumulation beyond the effect of ELF-PEMF alone. Figure 6A, presents the graphs for both inhibitors, depicting their respective effects on ROS levels.

**Fig. 6.**
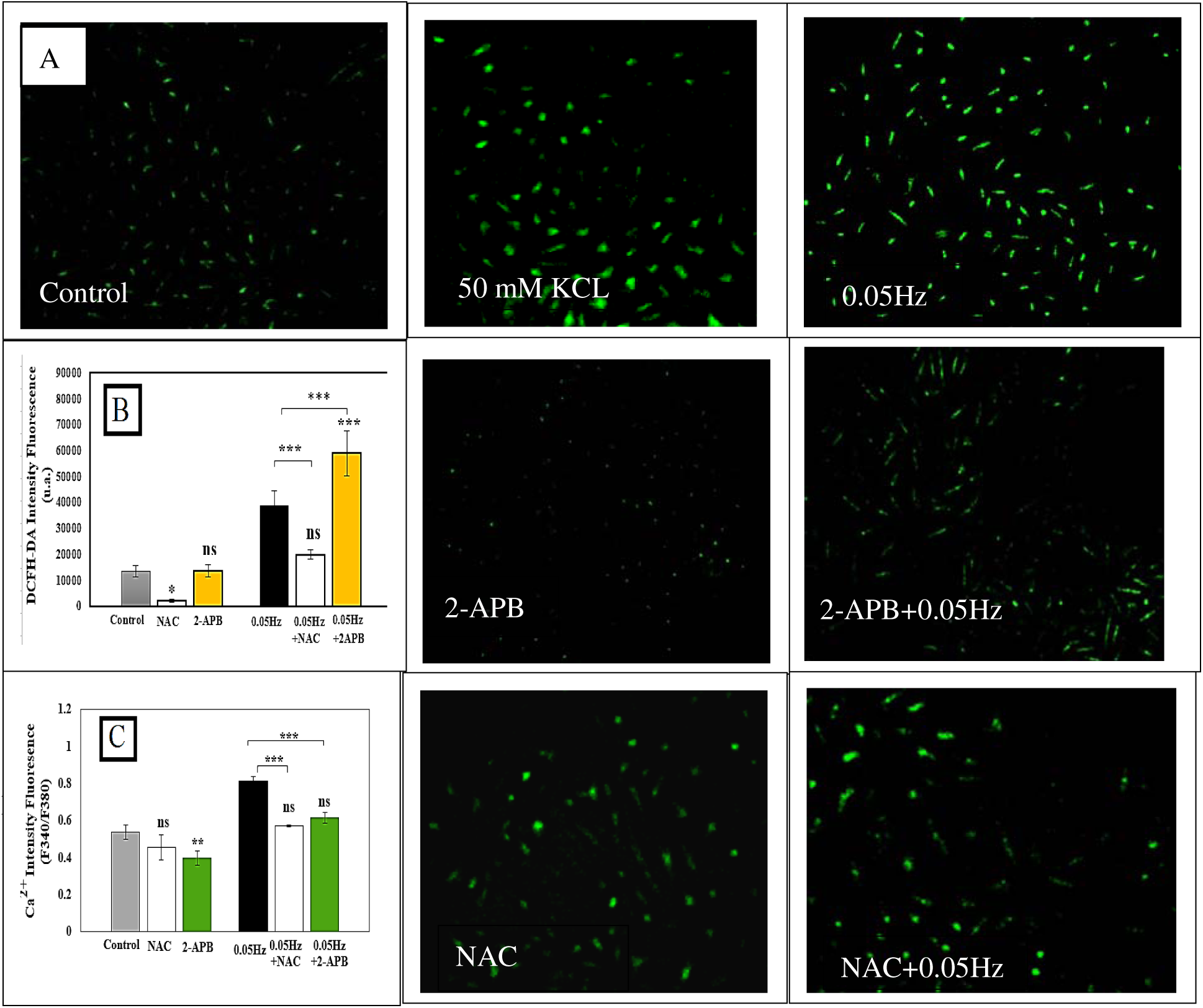
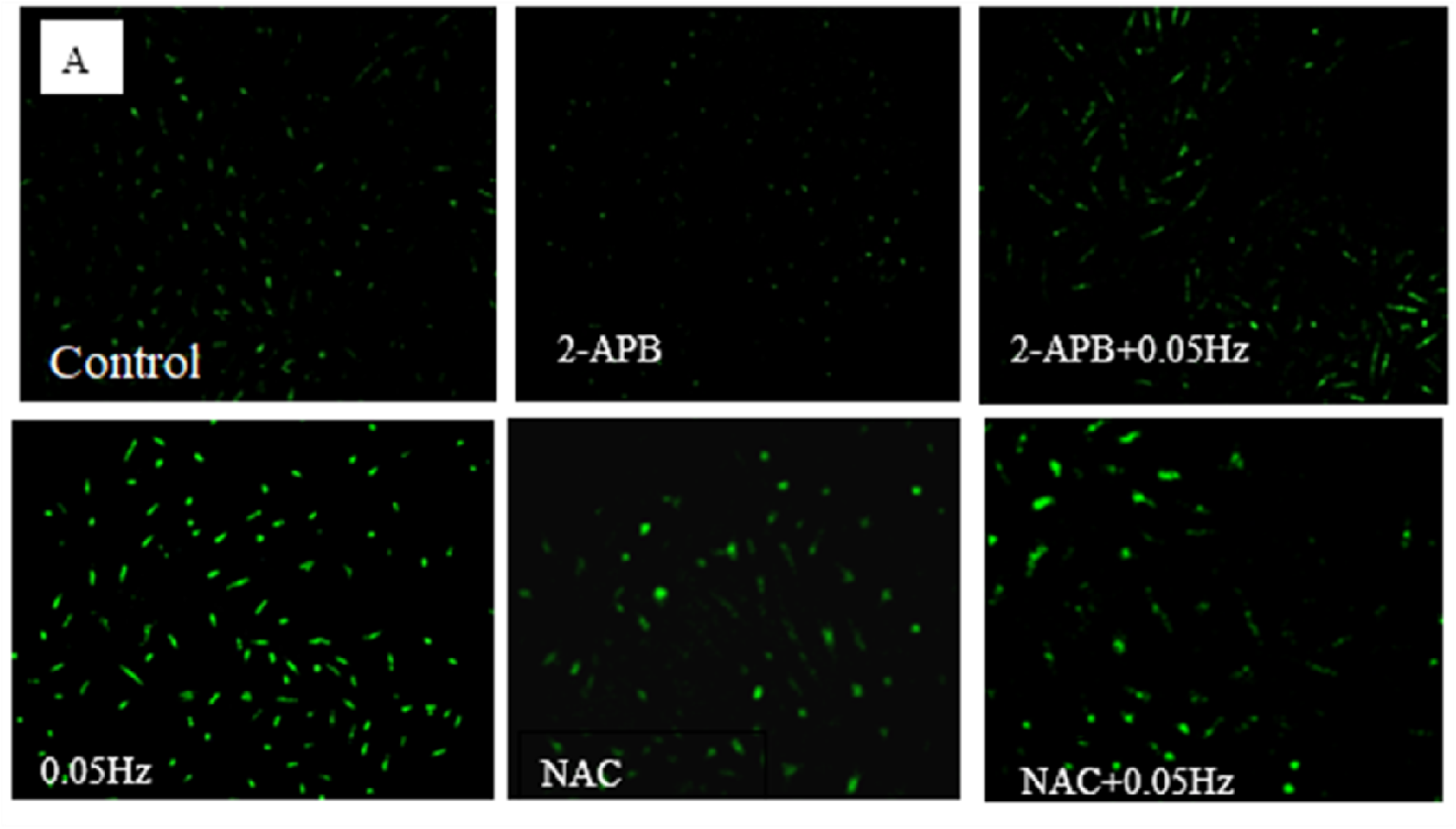
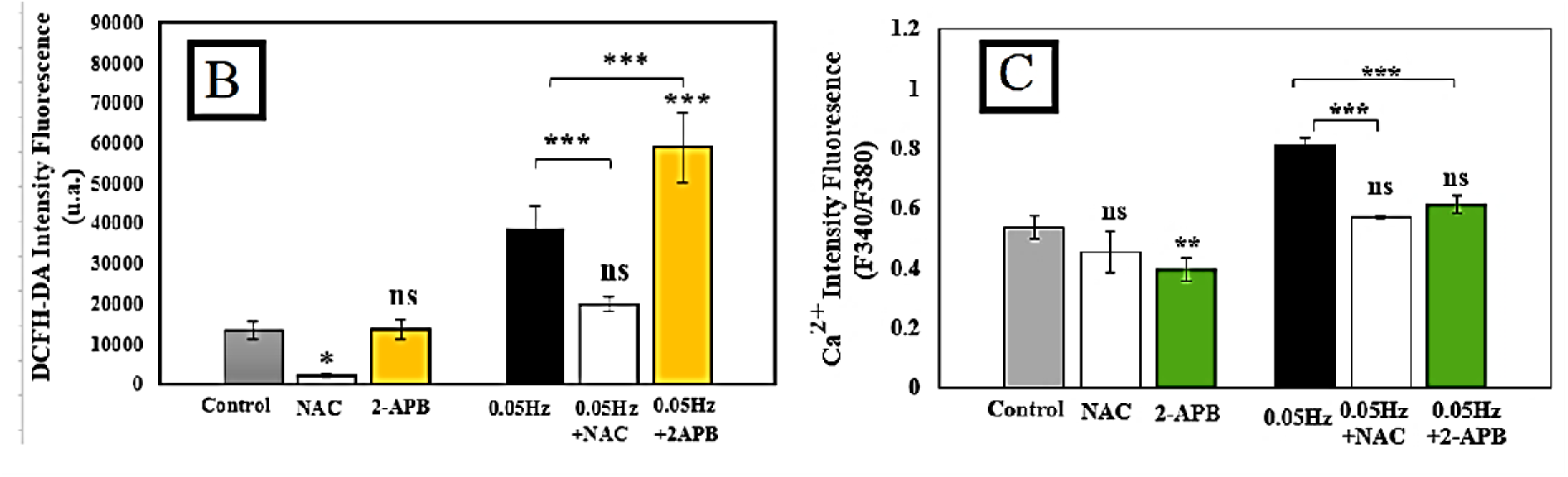
(A) Representative calcium fluorescence images (7 images) for control, KCl (50 mM), 0.05-Hz ELF-PEMF (100 mT, 45 min), and NAC- or 2-APB–treated groups with or without ELF-PEMF. (B) ROS quantification and (C) intracellular calcium responses under the same conditions. Data are mean ± SEM; one-way ANOVA with Tukey’s post hoc test: **p < 0.01, \*\*\**p < 0.0001, ns = not significant*.

**Fig. 7.**
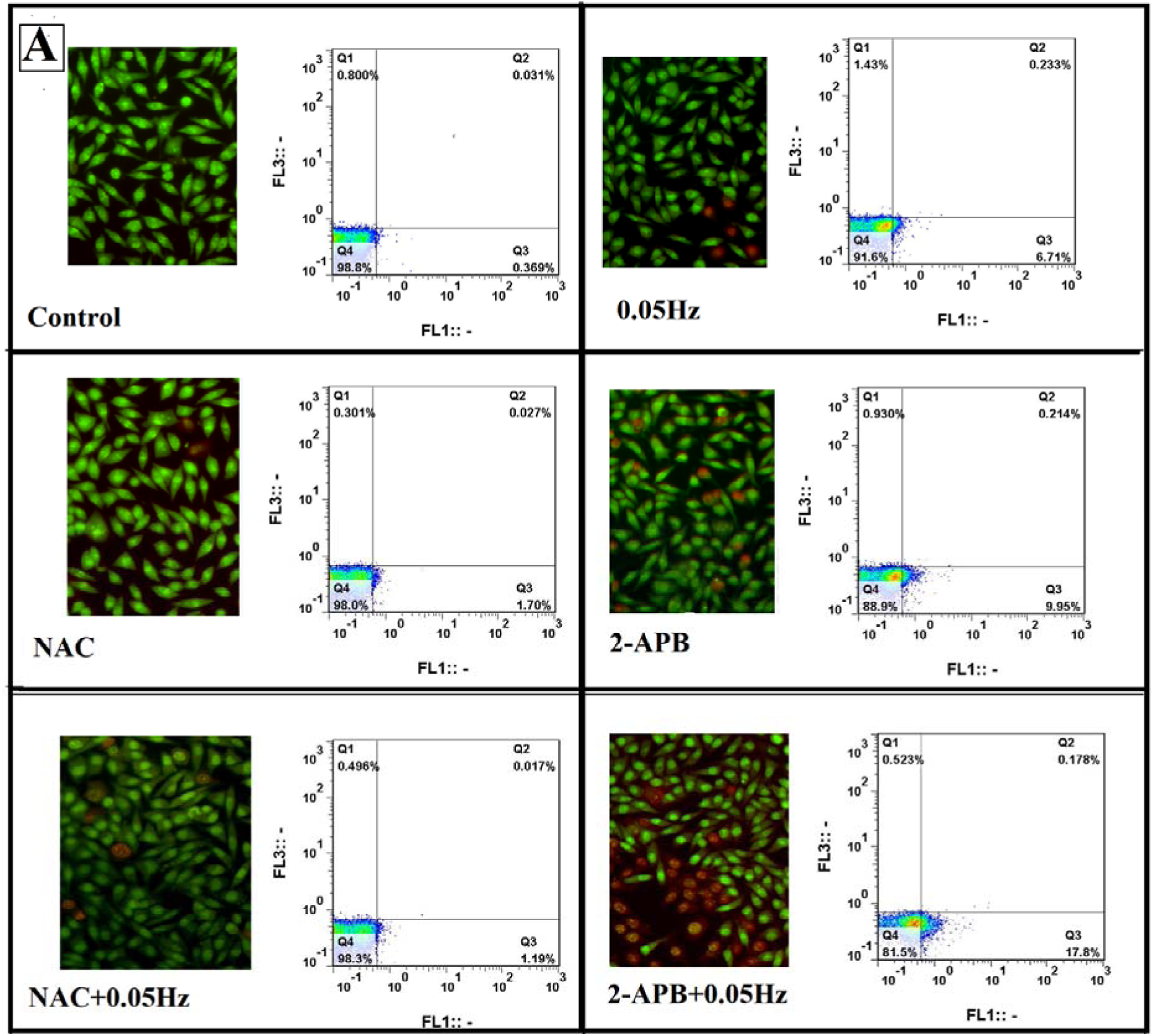
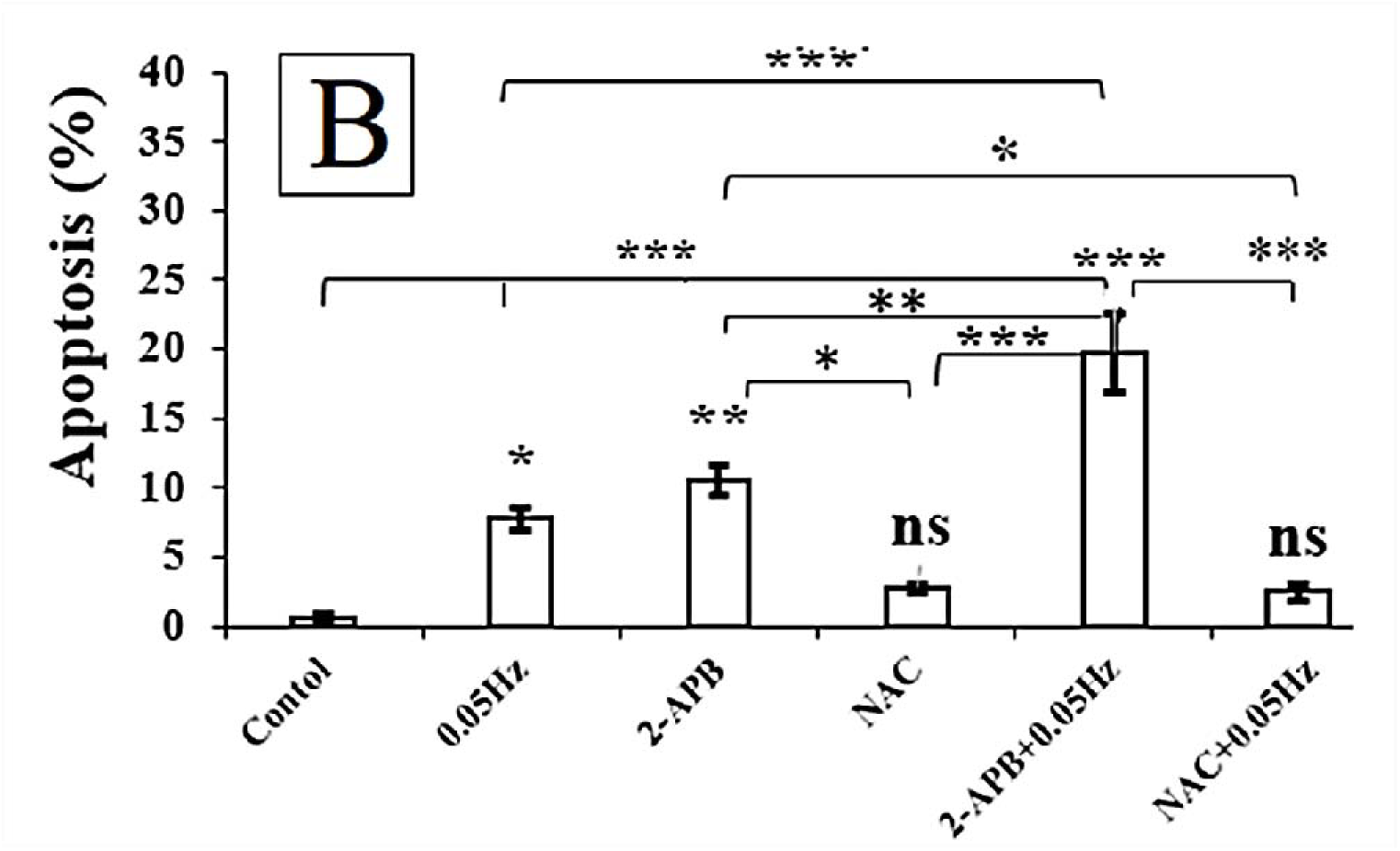
Microscopic images of apoptosis AO/EB staining of u87 cell death along with flow cytometry data for control, 0.05-Hz, NAC and 2-APB–treated with or without ELF-PEMF (100 mT, 45 min). Quantification of apoptotic cells (%), based on AO/EB. Data are mean ± SEM; significance determined by one-way ANOVA with Tukey’s test (*p < 0.05 vs. control) and other comparisons are only significant if they are shown.

#### 3.4.2 NAC and 2-APB Effects on Ca²D signaling

For the assessment of relative intracellular calcium intensity (F340/F380), U87 cells were again treated with 2-APB, which, as expected, significantly reduced the relative intracellular Ca²L intensity. However, when cells were simultaneously exposed to the 0.05 Hz ELF-PEMF field and 2-APB, the relative Ca²L intensity showed a slight, non-significant increase compared with the control, and no notable change was observed. In contrast, cells exposed to the ELF-PEMF field alone, as reported in previous sections, exhibited a significant increase in relative calcium intensity. Similarly, the relative intracellular Ca²L intensity in U87 cells treated with NAC was reduced compared to the control, but this reduction did not result in a significant change in the relative calcium intensity. Even when NAC treatment was combined with ELF-PEMF, despite a slight increase in the relative Ca²L intensity, this increase was not statistically compared to the control. All quantitative data are presented in Figure 6B.

### 3.5 Apoptosis Analysis

Apoptotic cell death was evaluated using acridine orange/ethidium bromide (AO/EB) staining in U87 glioblastoma cells exposed to ELF-PEMF at 0.05 Hz. In the absence of field exposure, apoptotic cells accounted for approximately 0.5 % in control samples, 1.7 % following NAC treatment, and 9.95% after 2-APB treatment. Upon ELF-PEMF exposure, apoptosis increased to 6.67% in 0.05Hz, 2.5% in NAC-treated cells and19.75 % in 2-APB-treated cells, indicating a differential modulation of cell death under field conditions. Flow cytometry analysis was performed as an independent quantitative method to validate the AO/EB findings. The flow cytometric data showed trends consistent with AO/EB staining, confirming the relative changes in apoptotic cell populations across experimental conditions.

## 4 Discussion

The biological effects of extremely ELF-PEMFs on cancer cells are strongly dependent on both intrinsic cellular characteristics and exposure parameters, including field intensity, frequency, and duration. In this study, U87 glioblastoma cells exhibited the most pronounced response at 0.05 Hz and 100 mT, consistent with a frequency-dependent “biological window” for optimal modulation of intracellular calcium (Ca²L) dynamics (24)To detect and quantify frequency and amplitude features of Ca²L oscillations, we applied FFT analysis (25), For samples exposed to ELF-PEMF, low-frequencies Ca²L oscillations exhibited higher amplitude, power, and energy compared with controls. Although these metrics decreased at higher frequencies, they remained elevated relative to controls, indicating sustained modulation of Ca²L signaling that may be associated with alterations in intracellular calcium homeostasis following ELF-PEMF exposure (26, 27). Furthermore, assessment of mitochondrial membrane potential (ΔΨm) in U87 cells following ELF-PEMF exposure revealed a decrease The associated significant increase in mitochondrial ROS indicates that, under these conditions, mitochondrial signaling may exceed adaptive thresholds and contribute to cellular stress and damage, likely mediated by transient intracellular Ca²L elevations(28, 29). Pharmacological intervention experiments in this study demonstrated that ROS production occurs upstream or independently of major Ca²L influx pathways.

Intracellular Ca²L release inhibition with 2-APB reduced cytosolic Ca²L below basal levels but did not prevent ROS accumulation, whereas antioxidant treatment with NAC effectively lowered ROS while maintaining basal Ca²L, highlighting that ROS modulation can occur independently of changes in intracellular Ca²L (30). These observations suggest that ELF-PEMF-induced oxidative stress is a primary event, with subsequent ROS–Ca²L crosstalk likely contributing to secondary alterations in calcium homeostasis. Ca²L oscillations in U87 cells are largely governed by ER release, SOCE, and mitochondrial buffering, with VGCCs likely playing a minimal role in ELF-PEMF responses. 2-APB reduces cytosolic Ca²L, highlighting the dominant contribution of IPL-dependent ER release and SOCE in both basal and ELF-PEMF-induced signaling(28). It is already shown that in the hippocampus, low-frequency ELF-EMF may modulate LTP via intracellular Ca²L release, an effect blocked by 2-APB and largely independent of VGCCs (31). Moreover, ELF-PEMF can influence cancer cells via mitochondrial radical pairs, amplifying ROS dynamics and altering oxidative stress. ELF-PEMF exposure appears to elevate ROS, likely through mitochondrial electron and proton (HL) leak, NADPH oxidase (NOX) activation, cryptochrome-mediated radical pair reactions, and local pH microdomains near mitochondria, NOX complexes, and ER, which may in turn influence ER Ca²L release, mitochondrial uptake, and SOCE (1, 32, 33). Notably, 2-APB may also contribute to some of these processes (2, 34). Consistently, in U87 cells, calcium inhibition with 2-APB did not fully prevent detectable cell death suggesting that ROS elevation may occur upstream and largely independently of Ca²L signaling. These Ca²L changes could subsequently engage downstream pathways, including MAPK/ERK and PI3K/Akt, potentially regulating proliferation, migration, survival, and stress adaptation in glioblastoma cells, although the precise molecular mechanisms remain to be fully elucidated (35, 36). The results of the present study, in agreement with previous reports, demonstrate that ELF-PEMFs can modulate intracellular calcium (Ca²L) levels and ROS production in glioma cells, highlighting these pathways as plausible mediators of their biological and therapeutic effects (14). Notably, analysis of our experimental data indicates that ROS elevation occurs prior to the increase in intracellular Ca²L, suggesting a potential upstream role for ROS in ELF-PEMF–induced cellular responses. Supporting this view, previous studies have demonstrated that ELF-PEMF exposure rapidly elevates intracellular ROS via cryptochrome and mitochondrial pathways, preceding significant changes in Ca²L dynamics or mitochondrial membrane potential, consistent with ROS acting as an early mediator of cellular responses (1, 4, 37). Studies on radiofrequency fields (e.g., 1.8 GHz) have shown direct modulation of ROS in human cells. While RF and ELF-PEMF differ in frequency and mechanism, these findings support the idea that electromagnetic fields can influence cellular redox balance, suggesting ROS may mediate ELF-PEMF effects (38) Given the well-established contribution of intracellular ROS accumulation to cancer cell death (39), these findings support the potential relevance of ELF-PEMFs as a therapeutic approach for glioma. Nevertheless, the precise molecular mechanisms linking ELF-PEMF exposure to ROS generation and subsequent Ca²L modulation remain insufficiently defined and warrant further investigation. Furthermore, the biological consequences of ELF-PEMF–induced ROS elevation are likely to be context-dependent, potentially involving alterations in redox-sensitive signaling pathways, disruption of cellular homeostasis, or activation of stress response and repair mechanisms (14, 40, 41). Importantly, the strong dependence of these effects on exposure parameters, including field intensity, amplitude, frequency, and duration as well as experimental and environmental conditions constitutes a key limitation of the present study and underscores the necessity for more advanced and standardized exposure platforms (42).

### Future research directions should focus on the following priorities

1. **Elucidation of early molecular events:** Detailed characterization of redox-and calcium-associated proteins that respond rapidly to ROS elevation, including redox-sensitive kinases, ion channels, and antioxidant response regulators, to define the initial signaling events triggered by ELF-PEMF exposure.
2. **Implementation of advanced biophysical models:** Adoption of modern quantum-informed and multiscale modeling frameworks to more accurately describe ELF-PEMF interactions with intracellular redox balance and calcium signaling networks.
3. **Integration of single-cell and systems-level methodologies:** Utilization of live-cell imaging, real-time redox and calcium sensors, and single-cell omics approaches—consistent with emerging Nature-class methodologies to capture cellular heterogeneity, strengthen causal inference, and identify molecular determinants of ELF-PEMF responsiveness.
4. **Evaluation of reproducibility and off-target effects:** Investigation of potential off-target effects and reproducibility across different glioma lines is warranted to ensure generalizability of findings

## 5 Conclusion

Results from this study indicate that ELF-PEMF exposure rapidly elevates intracellular ROS in U87 glioblastoma cells. Based on the observed patterns and existing literature, this increase is likely associated with mitochondrial electron/proton leak, activation of NADPH oxidase, and potentially cryptochrome-mediated radical pair mechanisms; however, these pathways were not directly dissected in the present work. ROS generation appeared to be independent of IPL-dependent ER Ca²L release, as inhibition with 2-APB did not prevent ROS elevation. The elevated ROS subsequently modulated downstream calcium signaling. Notably, the magnitude of these effects depended strongly on field frequency and amplitude, consistent with the concept of a frequency-dependent biological window. Collectively, these findings suggest that ROS may function as a primary mediator of ELF-PEMF-induced bioeffects in U87 cells, underscoring their potential relevance as a therapeutic target in glioblastoma. Nevertheless, further mechanistic studies are required to elucidate and confirm the underlying pathways.

## 6 Acknowledgments

The authors would like to express their gratitude to the Institute of Biochemistry and Biophysics (IBB), University of Tehran, and the Department of Biochemistry at Tarbiat Modares University for providing the necessary facilities and laboratory space for this research. We also thank the technical staff of the Molecular Biophysics Lab for their assistance during the ELF-PEMF exposure experiments and flow cytometry analysis.

## References

1. Sherrard RM, Morellini N, Jourdan N, El-Esawi M, Arthaut L-D, Niessner C, et al. Low-intensity electromagnetic fields induce human cryptochrome to modulate intracellular reactive oxygen species. PLoS biology. 2018;16(10):e2006229.

2. Zandieh A, Shariatpanahi SP, Ravassipour AA, Azadipour J, Nezamtaheri MS, Habibi-Kelishomi Z, et al. An amplification mechanism for weak ELF magnetic fields quantum-bio effects in cancer cells. Scientific Reports. 2025;15(1):2964.

3. Akdag MZ, Dasdag S, Cakir DU, Yokus B, Kizil G, Kizil M. Do 100-and 500-μT ELF magnetic fields alter beta-amyloid protein, protein carbonyl and malondialdehyde in rat brains? Electromagnetic Biology and Medicine. 2013;32(3):363–72.

4. Morabito C, Rovetta F, Bizzarri M, Mazzoleni G, Fanò G, Mariggiò MA. Modulation of redox status and calcium handling by extremely low frequency electromagnetic fields in C2C12 muscle cells: A real-time, single-cell approach. Free Radical Biology and Medicine. 2010;48(4):579–89.

5. Chen Y. Exposure to extremely low-frequency pulsed electromagnetic fields promotes fracture healing through alleviating the damage caused by cigarette smoke and modulating the osteoimmune microenvironment: Eberhard Karls Universität Tübingen; 2023.

6. Jimenez-Del-Rio M, Velez-Pardo C. The bad, the good, and the ugly about oxidative stress. Oxidative medicine and cellular longevity. 2012;2012(1):163913.

7. Palmeira CM, Teodoro JS, Amorim JA, Steegborn C, Sinclair DA, Rolo AP. Mitohormesis and metabolic health: The interplay between ROS, cAMP and sirtuins. Free Radical Biology and Medicine. 2019;141:483–91.

8. Katona M, Bartók Á, Nichtova Z, Csordás G, Berezhnaya E, Weaver D, et al. Capture at the ER-mitochondrial contacts licenses IP3 receptors to stimulate local Ca2+ transfer and oxidative metabolism. Nature communications. 2022;13(1):6779.

9. Feissner RF, Skalska J, Gaum WE, Sheu S-S. Crosstalk signaling between mitochondrial Ca2+ and ROS. Frontiers in bioscience: a journal and virtual library. 2009;14:1197.

10. Berridge MJ, Bootman MD, Roderick HL. Calcium signalling: dynamics, homeostasis and remodelling. Nature reviews Molecular cell biology. 2003;4(7):517–29.

11. Reane DV, Rizzuto R, Raffaello A. The ER-mitochondria tether at the hub of Ca2+ signaling. Current Opinion in Physiology. 2020;17:261–8.

12. Smedler E, Uhlén P. Frequency decoding of calcium oscillations. Biochimica et Biophysica Acta (BBA)-General Subjects. 2014;1840(3):964–9.

13. Barhoumi R, Qian Y, Burghardt RC, Tiffany-Castiglioni E. Image analysis of Ca2+ signals as a basis for neurotoxicity assays: promises and challenges. Neurotoxicology and teratology. 2010;32(1):16–24.

14. Huang M, Li P, Chen F, Cai Z, Yang S, Zheng X, et al. Is extremely low frequency pulsed electromagnetic fields applicable to gliomas? A literature review of the underlying mechanisms and application of extremely low frequency pulsed electromagnetic fields. Cancer medicine. 2023;12(3):2187–98.

15. Delierneux C, Kouba S, Shanmughapriya S, Potier-Cartereau M, Trebak M, Hempel N. Mitochondrial calcium regulation of redox signaling in cancer. Cells. 2020;9(2):432.

16. Kang SS, Han K-S, Ku BM, Lee YK, Hong J, Shin HY, et al. Inhibition of the Ca2+ release channel, IP3R subtype 3 by caffeine slows glioblastoma invasion and migration and extends survival. Cancer Research. 2010;70(3):1173.

17. Maklad A, Sharma A, Azimi I. Calcium signaling in brain cancers: roles and therapeutic targeting. Cancers. 2019;11(2):145.

18. Nezamtaheri MS, Goliaei B, Shariatpanahi SP, Ansari AM. Differential biological responses of adherent and non-adherent (cancer and non-cancerous) cells to variable extremely low frequency magnetic fields. Scientific Reports. 2022;12(1):14225.

19. Grynkiewicz G, Poenie M, Tsien RY. A new generation of Ca2+ indicators with greatly improved fluorescence properties. Journal of biological chemistry. 1985;260(6):3440–50.

20. Abcam. Fura-2 AM calcium imaging protocol 2026 [updated January 30, 2026. Available from: https://www.abcam.com/en-us/technical-resources/protocols/fura-2-am-imaging.

21. DeHaven WI, Smyth JT, Boyles RR, Bird GS, Putney Jr JW. Complex actions of 2-aminoethyldiphenyl borate on store-operated calcium entry. Journal of Biological Chemistry. 2008;283(28):19265–73.

22. Xu S-Z, Zeng F, Boulay G, Grimm C, Harteneck C, Beech DJ. Block of TRPC5 channels by 2-aminoethoxydiphenyl borate: a differential, extracellular and voltage-dependent effect. British journal of pharmacology. 2005;145(4):405.

23. Zhitkovich A. N-acetylcysteine: antioxidant, aldehyde scavenger, and more. ACS Publications; 2019. p. 1318–9.

24. López de Mingo I, Rivera González M-X, Ramos Gómez M, Maestú Unturbe C. The Frequency of a Magnetic Field Determines the Behavior of Tumor and Non-Tumor Nerve Cell Models. International Journal of Molecular Sciences. 2025;26(5):2032.

25. Skupin A, Falcke M. Statistical analysis of calcium oscillations. The European Physical Journal Special Topics. 2010;187(1):231–40.

26. Golbach LA, Portelli LA, Savelkoul HF, Terwel SR, Kuster N, de Vries RB, et al. Calcium homeostasis and low-frequency magnetic and electric field exposure: A systematic review and meta-analysis of in vitro studies. Environment international. 2016;92:695–706.

27. Zhao Y-L, Yang J-C, Zhang Y-H, editors. Effects of magnetic fields on intracellular calcium oscillations. 2008 30th Annual International Conference of the Ieee Engineering in Medicine and Biology Society; 2008: IEEE.

28. Luo F-L, Yang N, He C, Li H-L, Li C, Chen F, et al. Exposure to extremely low frequency electromagnetic fields alters the calcium dynamics of cultured entorhinal cortex neurons. Environmental research. 2014;135:236–46.

29. Peng B, Wang Y, Zhang H. Mitonuclear Communication in Stem Cell Function. Cell Proliferation. 2025;58(5):e13796.

30. Vezir Ö, Çömelekoğlu Ü, Sucu N, Yalın AE, Yılmaz ŞN, Yalın S, et al. N-Acetylcysteine-induced vasodilatation is modulated by KATP channels, Na+/K+-ATPase activity and intracellular calcium concentration: An in vitro study. Pharmacological Reports. 2017;69(4):738–45.

31. Zhao W, Dong L, Tian L, Zhao L, Zhao Y, Zheng Y. Changes in intracellular calcium concentration level accompany ageLrelated inhibitions of longLterm potentiation in hippocampus induced by extremely low frequency electromagnetic fields. European Journal of Neuroscience. 2023;58(2):2437–50.

32. Georgiou CD, Margaritis LH. Oxidative stress and NADPH oxidase: connecting electromagnetic fields, cation channels and biological effects. International Journal of Molecular Sciences. 2021;22(18):10041.

33. Teranishi M, Ito M, Huang Z, Nishiyama Y, Masuda A, Mino H, et al. Extremely low-frequency electromagnetic field (ELF-EMF) increases mitochondrial electron transport chain activities and ameliorates depressive behaviors in mice. International Journal of Molecular Sciences. 2024;25(20):11315.

34. Dubinin MV, Chulkov AV, Igoshkina AD, Cherepanova AA, Mikina NV. Effect of 2-aminoethoxydiphenyl borate on the functions of mouse skeletal muscle mitochondria. Biochemical and Biophysical Research Communications. 2024;712:149944.

35. Vitucci M, Karpinich NO, Bash RE, Werneke AM, Schmid RS, White KK, et al. Cooperativity between MAPK and PI3K signaling activation is required for glioblastoma pathogenesis. Neuro-oncology. 2013;15(10):1317–29.

36. Lai Y, Lu X, Liao Y, Ouyang P, Wang H, Zhang X, et al. Crosstalk between glioblastoma and tumor microenvironment drives proneural–mesenchymal transition through ligand-receptor interactions. Genes & Diseases. 2024;11(2):874–89.

37. Wang H, Zhang X. Magnetic fields and reactive oxygen species. International journal of molecular sciences. 2017;18(10):2175.

38. Pooam M, Jourdan N, Aguida B, Dahon C, Baouz S, Terry C, et al. Exposure to 1.8 GHz radiofrequency field modulates ROS in human HEK293 cells as a function of signal amplitude. Communicative & Integrative Biology. 2022;15(1):54–66.

39. An X, Yu W, Liu J, Tang D, Yang L, Chen X. Oxidative cell death in cancer: mechanisms and therapeutic opportunities. Cell death & disease. 2024;15(8):556.

40. Gorrini C, Harris IS, Mak TW. Modulation of oxidative stress as an anticancer strategy. Nature reviews Drug discovery. 2013;12(12):931–47.

41. Trachootham D, Alexandre J, Huang P. Targeting cancer cells by ROS-mediated mechanisms: a radical therapeutic approach? Nature reviews Drug discovery. 2009;8(7):579–91.

42. Mansourian M, Shanei A. Evaluation of pulsed electromagnetic field effects: a systematic review and metaLanalysis on highlights of two decades of research in vitro studies. BioMed Research International. 2021;2021(1):6647497.

